# Characterizing the structure-function relationship of a naturally-occurring RNA thermometer

**DOI:** 10.1101/142141

**Authors:** Sarai Meyer, Julius B. Lucks

## Abstract

A wide number of bacteria have been found to govern virulence and heat shock responses using temperature-sensing RNAs known as RNA thermometers. A prime example is the *agsA* thermometer known to regulate the production of the AgsA heat shock protein in *Salmonella enterica* using a “fourU” structural motif. Using the SHAPE-Seq RNA structure-probing method *in vivo* and *in vitro*, we found that the regulator functions by a subtle shift in equilibrium RNA structure populations that lead to a partial melting of the helix containing the ribosome binding site. We also demonstrate that ribosome binding to the *agsA* mRNA causes changes to the thermometer structure that appear to facilitate thermometer helix unwinding. These results demonstrate how subtle RNA structural changes can govern gene expression and illuminate the function of an important bacterial regulatory motif.

---

Bacterial survival depends on a bacterium’s ability to react appropriately to environmental cues, altering its behavior in response to its ever-changing surroundings. In particular, variations in temperature require coordinated system-wide responses in order to avoid deleterious effects,^1^ resulting in the evolution of a wide array of molecular thermosensors. Among these thermosensors, RNA thermometers have been shown to control gene expression in response to changes in external temperature, allowing bacteria to govern wide-spread responses to heat shock and to control temperature-dependent virulence effects,^2^ an important feature for pathogens with warm-blooded hosts.

RNA thermometers generally govern heat shock response post-transcriptionally by relying on the formation of temperature-sensitive secondary structures to fully or partially occlude the ribosome binding site (RBS) of the controlled gene,^3,4^ thus decreasing translation efficiency^5^ at low temperatures. An elevation in temperature changes this structure and allows translation in a temperature-dependent manner. The most widely characterized RNA thermometers are the broad family of ROSE (<u>r</u>epression <u>o</u>f heat <u>s</u>hock gene <u>e</u>xpression) elements common among α- and γ-proteobacteria^6,7^ and found in the 5’ untranslated regions (UTRs) of heat shock genes.^1^ ROSE elements share a conserved sequence motif that sequesters the Shine-Dalgarno (SD) sequence in a hairpin through an unusual base triple and non-canonical nucleotide interactions.^8^

Other heat shock RNA thermometers involve the formation of complex structures that block ribosome entry without directly pairing with the RBS,^9,10^ or depend upon an internal hairpin bulge to allow for stem melting in response to increased temperature.^11,12^ This hairpin bulge pattern occurs in the *Salmonella enterica agsA* RNA thermometer (Figure 1A), where an internal stem bulge is followed by a stretch of four uridines paired to the Shine-Dalgarno sequence. This structure confers thermosensing capability to the 5’ UTR of the *agsA* gene encoding the aggregation suppression protein AgsA,^13^ a molecular chaperone that forms part of the heat shock response in *Salmonella*.^14^ Although the fourU RNA thermometer motif was first coined in reference to the *agsA* thermometer from *Salmonella*,^13^ the previously reported virulence control mechanism of the plague bacterium *Yersinia pestis* also relies on a four uridine base-pairing to the Shine-Dalgarno sequence to grant temperature-dependent translational control.^15^ Similarly, the temperature-dependent virulence response of *Vibrio cholerae* was recently tied to a fourU element within the 5’ UTR of ToxT, a key transcriptional activator of virulence factors.^16^ While other temperature-based virulence controls rely on alternate mechanisms,^17,18^ the reoccurrence of the fourU motif in regulating both heat shock and virulence underlines its widespread utility as an RNA thermometer.

**Figure 1.**
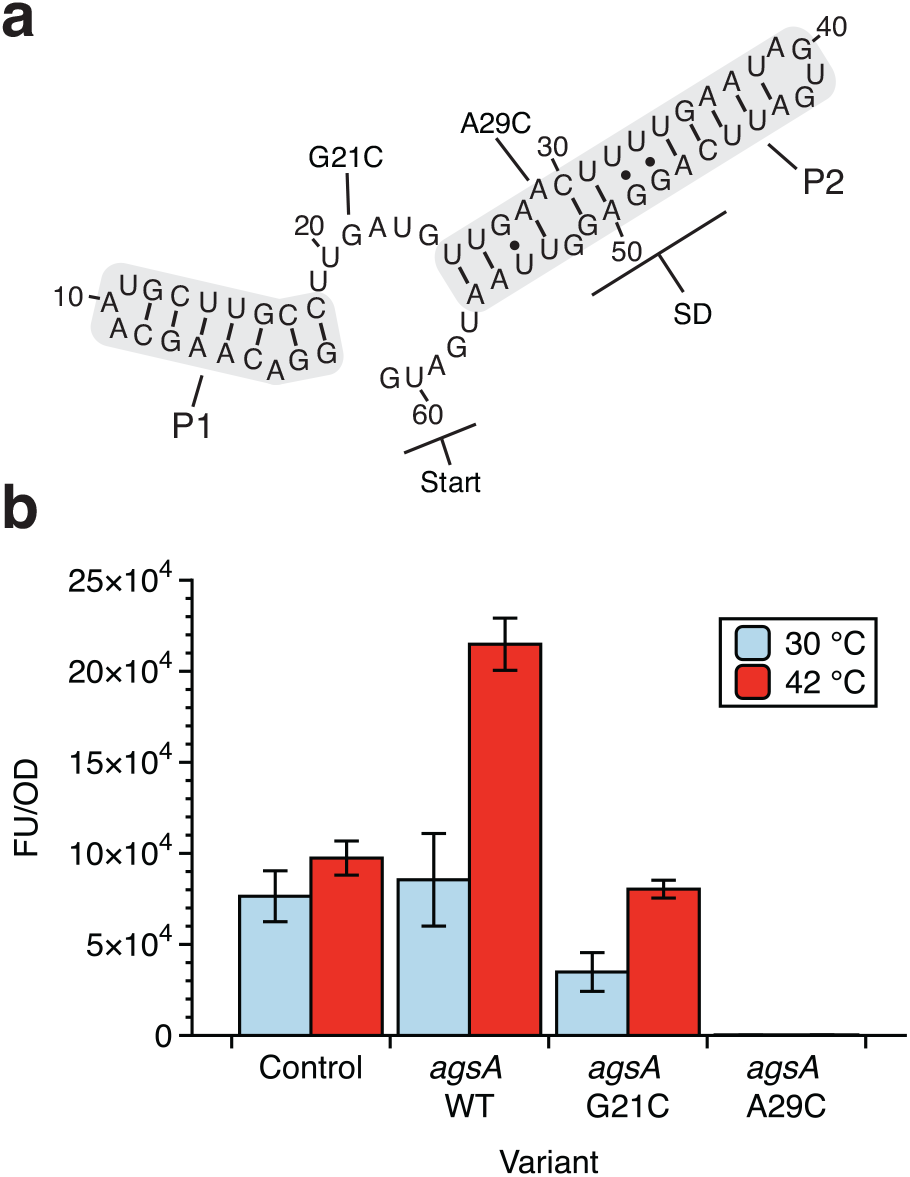
Structure and function of the *agsA* thermometer. (**a**) Putative structure of the *agsA* 5’ untranslated region (UTR) RNA thermometer from *Salmonella enterica* with key P1 and P2 helices, the Shine-Dalgarno (SD) sequence, start codon, and mutations of interest from Waldminghaus et al.^13^ labeled. (**b**) Functional *in vivo* testing of the *agsA* thermometer and mutants in *E. coli* using the superfolder green fluorescent protein (SFGFP) reporter gene placed downstream of the 5’ UTR. Cultures were grown at 30 °C for four hours, then subjected to 30 minutes of 30 °C or 42 °C to measure the response of the thermometer constructs to heat shock. The G21C mutation extends the P2 helix to include the start codon, resulting in lower fluorescence in both the low temperature and heat shock conditions. The A29C mutation closes the inner-hairpin loop within the P2 helix, causing complete repression of gene expression under both temperature conditions. Fluorescence is normalized as fluorescence units over optical density (FU/OD).

The fourU motif hairpin is intuitively thought to function by simply opening at elevated temperatures to allow ribosome access, though it is unclear exactly how ‘open’ the structure must be to implement temperature-dependent control. In fact, *agsA* thermometer structural characterization *in vitro* via NMR spectroscopy has given key insights into the relative stabilities of the different pairings within the thermometer hairpin^19^ and the magnesium-dependence of hairpin melting.^20^ This previous work has also shown that only a few internal bases have increased flexibility at elevated temperatures, rather than large structural changes indicating melting of the entire helix.

Here we sought to better understand the mechanistic details of the *agsA* fourU thermometer by studying for the first time how *in vivo* structural changes couple to gene expression. By combining in-cell SHAPE-Seq to probe RNA structures *in vivo*, fluorescent reporter measurements of gene expression, and experimentally restrained computational structure prediction, we present a detailed view of the *agsA* thermometer structural ensemble and its response to heat shock. We then dissect the underlying structural mechanism through *in vitro* experiments performed in cell free gene expression systems and highlight the effect of ribosome binding and translation on the structure of the *agsA* 5’ UTR. This detailed study of the *agsA* thermometer illuminates the intriguing relationship between subtle changes in its RNA structure and large changes in its regulatory function.

## Experimental Procedures

### Cloning and plasmid construction

The *agsA* 5’ UTR sequence from Waldminghaus et al.^13^ in both its wild-type (WT) and G21C and A29C point-mutant forms was cloned into a p15a plasmid backbone harboring chloramphenicol resistance. Using inverse PCR (iPCR) for scarless insertion, the *agsA* sequence was introduced directly between the *Escherichia coli* sigma 70 consensus promoter J23119 SpeI (a variant of the BBa_J23119 promoter from the Registry of Standard Biological Parts http://partsregistry.org/Main_Page in which the final six nucleotides have been replaced with a SpeI cut site) and the start codon for the superfolder green fluorescent protein (SFGFP) reporter gene. The constitutive gene expression control plasmid has a sequence designed to allow full ribosome access to the RBS. See Supplementary Information for plasmid maps (Figure S1) and sequences (Table S1).

### In vivo bulk fluorescence assays and in-cell SHAPE-Seq

All bulk fluorescence and in-cell SHAPE-Seq experiments were performed in *E. coli* strain TG1 *(F’traD36 lacIq Delta(lacZ) M15 pro A+B+/supE Delta(hsdM-mcrB)5 (rk- mk- McrB-) thi Delta(lac-proAB))* with three independent biological replicates. The overall heat shock protocol for testing *agsA* responsiveness to heat shock is depicted in Figure S2. For each independent replicate, an empty control plasmid (JBL 001), constitutive gene expression control plasmid (JBL2348), and *agsA* WT and mutant plasmids were transformed separately into chemically competent *E. coli* TG1 cells, plated on LB + Agar plates with 34 mg/ml chloramphenicol and incubated overnight at 37 °C for approximately 17 hours. After overnight incubation, the plates were removed from the incubator and kept at room temperature for approximately 7 hours. For each of the three replicates, three separate colonies were picked for the control and one of each tested variant, and each colony was used to inoculate 1 mL of LB media with 34 mg/mL chloramphenicol in a 2 mL 96-well block. The block was covered with a breathable seal and incubated at 37 °C while shaking at a speed of 1,000 rpm in a bench-top shaker for 18 hours overnight. From these overnight cultures, 24 *μ*L were taken and used to inoculate two duplicate 96-well blocks containing 1.2 mL LB media with 34 mg/mL chloramphenicol. The blocks were shaken at 30 °C and 1,000 rpm for 4 hours, and then one of the blocks was moved to shake at 42 °C (heat shock condition), while the other was kept at 30 °C for 30 min (no heat shock condition). From there, in-cell SHAPE-Seq probing (for RNA structure determination) and bulk fluorescence sample preparation (for gene expression measurement) were carried out as described by Watters et al.^21,22^ using 1 mL of the sample for in-cell SHAPE-Seq and 150 *μ*L for the bulk fluorescence measurements. Any deviations from the published protocol are described below. The samples’ optical density (OD) at 600 nm and fluorescence (485 nm excitation, 520 nm emission) were measured with a Biotek Synergy H1 plate reader for the bulk fluorescence. Following RNA probing as described,^21^ the modified and unmodified RNAs were extracted using the TRIzol Max Bacterial RNA Isolation Kit (Thermo Fisher). The extracted RNAs were reverse transcribed from within the SFGFP gene (primer = CAACAAGAATTGGGACAACTCCAGTG) using 3 *μ*L of 50 nM (not 0.5 *μ*M) primer per reaction. The following RNA hydrolysis step was extended from 5 minutes to 15 minutes, and the partial neutralization with HCl was omitted. Subsequent ethanol precipitation, adaptor ligation, and bead purification steps followed the published protocol exactly,^21^ and the quality analysis and library construction were performed following the previously reported PCR modifications to minimize side-product formation for libraries from weakly expressed RNAs.^21^ Data analysis was executed using the Spats pipeline version 1.0.1 as described by Watters et al.^22^ in order to generate raw θ reactivities,^23^ which were then converted to ϱ reactivities by multiplying by the length of the probed RNA^24^ and normalized using the box-plot strategy described by Low and Weeks.^25^ The normalization method ignores outlier reactivities, defined as those reactivities more than 1.5 times the interquartile range (IQR) above the 75^th^ percentile, and then normalizes all reactivities by the average of the next 10 percent of reactivities. This normalization scheme is designed to allow for comparison of reactivities between structures with varying distributions of paired and unpaired nucleotides.

### mRNA preparation

DNA templates for *in vitro* transcription were generated using PCR to add the T7 promoter to the *agsA* variant and control sequences, and to amplify the region from the 5’ UTR to the intrinsic terminator following the SFGFP gene. The resulting templates were *in vitro* transcribed with T7 polymerase, and the resulting mRNAs were purified using Ampure XP beads (Beckman Coulter) following the manufacturer’s protocol and run alongside an RNA ladder on a 3% denaturing polyacrylamide gel. The gel was stained using SYBR Gold (Thermo Fisher) and visualized under ultraviolet light to check for template length and purity.

### PURExpress assays

The cell-free assays showing temperature-sensitive expression from both *agsA* DNA templates and pre-transcribed mRNA were performed in the PURExpress ΔRibosome Kit (New England Biolabs). To each transcription-translation reaction were added 4 *μ*L of kit Solution A, 1.2 uL Factor Mix, 1.8 uL *E. coli* ribosomes (New England Biolabs), 0.75 *μ*L *E. coli* RNA polymerase holoenzyme (New England Biolabs), 0.4 uL SUPERase In RNase Inhibitor (Thermo Fisher), 100 ng plasmid template, and enough nuclease-free water to reach a total volume of 10 *μ*L. Each translation-only reaction contained 4 *μ*L Solution A, 1.2 uL Factor Mix, 1.8 uL *E. coli* ribosomes, 3.24 pmol purified mRNA template, and enough nuclease-free water to reach a total volume of 10 *μ*L. Reactions were pipetted into a clear-bottomed, black 384-well plate loaded onto a Biotek Synergy H1 plate reader pre-warmed to either 30 °C or 42 °C where fluorescence (485 nm excitation, 520 nm emission) was monitored for 3 hours with measurements taken every 5 minutes. Measurements were corrected for background fluorescence by subtracting the average reading for three empty wells.

### In vitro SHAPE-Seq in folding buffer

*In vitro* SHAPE-Seq was carried out largely following the *in vitro* SHAPE-Seq protocol from Watters et al.^22^ performed on 10 pmol of purified mRNA folded and probed at either 30 °C or 42 °C in 18 *μ*L of folding buffer (final concentration: 10 mM MgCl_2_, 100 mM NaCl, and 100 mM HEPES). The folded RNA was split into two 9 *μ*L aliquots and probed with the addition of 1 *μ*L 65 mM 1M7 (1-methyl-7-nitroisatoic anhydride) in dimethyl sulfoxide (DMSO) (positive channel) or 1 *μ*L DMSO (negative channel). After modification, the RNAs were ethanol precipitated with the addition of 90 *μ*L nuclease-free water, 10 *μ*L 3 M sodium acetate (pH 5.5), 1 *μ*L glycogen, and 300 *μ*L ice-cold 100% ethanol, followed by freezing for 30 minutes at −80 °C and centrifugation at 4 °C and 15,000 rpm for 30 minutes. All liquid was aspirated, and the pellets were washed with 200 *μ*L 70% ethanol before further centrifugation for 2 minutes, removal of the wash liquid, and re-suspension in 10 *μ*L of nuclease-free water. From there, the remainder of the experimental protocol—from reverse transcription through raw reactivity calculation followed the published method from Watters et al.^22^ Reactivities were normalized as above.

### In vitro SHAPE-Seq in PURExpress

For experiments probing the structure of the *agsA* WT and control mRNAs in PURExpress without ribosomes, with active translation, or with stalled translation initiation complexes, 3.24 picomoles of purified mRNA were probed at either 30 °C or 42 °C. The translation initiation complex reactions were assembled using the PURExpress Δaa ΔtRNA kit (New England Biolabs), with each 10 *μ*L reaction containing 2 *μ*L kit Solution A, 1 *μ*L 3 mM L-methionine, 1 *μ*L kit tRNA, 3 *μ*L kit Solution B, and 3.24 picomoles of mRNA — the lack of amino acids other than methionine was designed to cause ribosomal complex stalling at the start codon. The PURExpress probing experiments without ribosomes were carried out in a similar fashion using the PURExpress ΔRibosome Kit (New England Biolabs), with each 10 *μ*L reaction containing 4 *μ*L kit Solution A, 1.2 *μ*L Factor Mix, and 3.24 picomoles of mRNA. The PURExpress active translation reactions were assembled identically except for the addition of 1.2 uL of *E. coli* ribosomes. PURExpress reactions were incubated for 20 minutes (no ribosomes and stalled translation initiation complexes) or 35 minutes (active translation) at 30 °C or 42 °C before being split for modification—5 *μ*L of reaction were added to 0.56 *μ*L 65 mM 1M7 in DMSO (positive channel) and to 0.56 *μ*L DMSO (negative channel) and incubated for an additional minute before being transferred to ice. From there, the modified and unmodified RNA were extracted using the Trizol Max Bacterial RNA Isolation Kit (Thermo Fisher), and the remainder of the protocol (including reactivity normalization) followed the same method as the in-cell SHAPE-Seq samples.

### Statistical analysis and data presentation

Experiments were carried out with three distinct replicates. Error bars represent sample standard deviation, and significance was determined using two-sided heteroscedastic Welch’s t-tests. For SHAPE-Seq difference plots, error was propagated using standard propagation of error techniques. Reactivity normalization was carried out as described in the in-cell SHAPE-Seq section above.

### RNAStructure structure prediction and MDS analysis

Minimum free energy (MFE) RNA structures were predicted using the `Fold` command from the RNAStructure command-line interface.^26^ Box-plot normalized reactivities were specified as pseudoenergy restraints using RNAStructure’s “`−sh`” option, and the SHAPE restraint intercept (*b*) and slope (*m*) were set at −0.3 kcal/mol and 1.1 kcal/mol, respectively, based on the optimum values for weighting ϱ reactivities as determined by Loughrey et al.^24^ Temperature was specified as either 303.15 K (30 °C) or 315.15K (42 °C) using the “`−t`” temperature option. Structures were output as connectivity (ct) files, converted to dot-bracket notation (dbn) using the ct2dot utility from RNAStructure, and drawn using Adobe Illustrator (Figures 2b, 3d, 5b, S7, and S9b) or the VARNA graphical user interface (Figures S3-S6).^27^ Structures in Figures 2b, 3d, 5b, S7, and S9b represent predictions for *agsA* variant 5’ UTRs. Structures including the full mRNA sequence restrained by SHAPE data for the 5’ UTR and SFGFP leader sequence are presented in Figures S3 through S6. Partition function generation for the RNA probability pairing matrices was performed using the RNAStructure webserver’s `partition` function, with the same temperature and pseudoenergy SHAPE restraints as the MFE structure prediction, and probability plots were generated using the `ProbabilityPlot` function. For multidimensional scaling (MDS) analysis of the RNA structural ensemble, 100,000 structures were sampled from each condition using the RNAStructure `stochastic` function. Following Kutchko et al.,^28^ all unique structures were coded as binary vectors based on whether each nucleotide was base-paired (0: not base-paired, 1: base-paired), and the resulting binary vectors were subjected to 2-component multidimensional scaling carried out with the R Project for Statistical Computing (v3.2.3).

**Figure 2.**
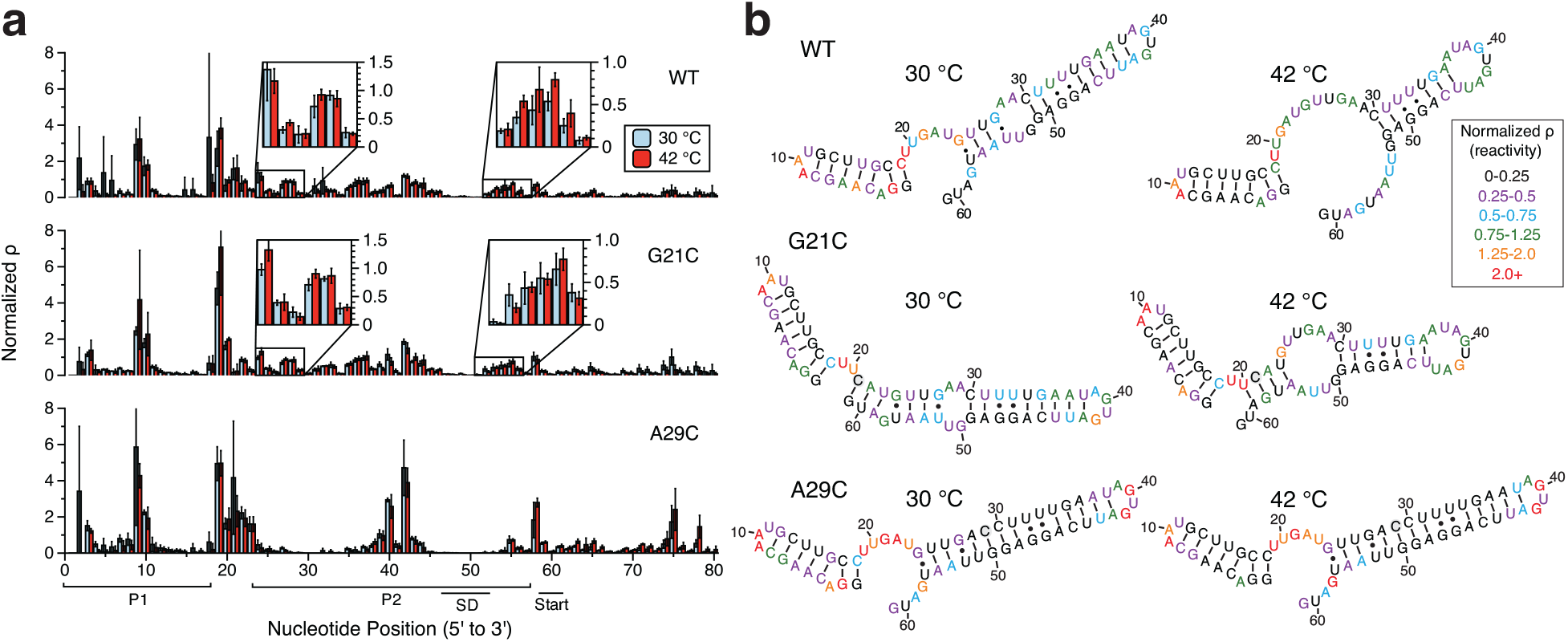
SHAPE-Seq characterization of the *agsA* thermometer *in vivo*. (**a**) SHAPE-Seq normalized reactivity (ϱ) plots for *agsA* WT, G21C, and A29C variants from triplicate experiments *in vivo* performed at 30 °C and using 42 °C heat shock conditions. Error bars represent standard deviations over replicates. (**b**) Experimentally-informed minimum free energy (MFE) RNA structures as predicted using the RNAStructure Fold program with the reactivity values from (a) taken as pseudoenergy folding restraints.^26^ Nucleotides are color-coded according to normalized SHAPE reactivity (ϱ), with red indicating high reactivity and black low reactivity. Point-mutated nucleotides are shaded in grey.

**Figure 3.**
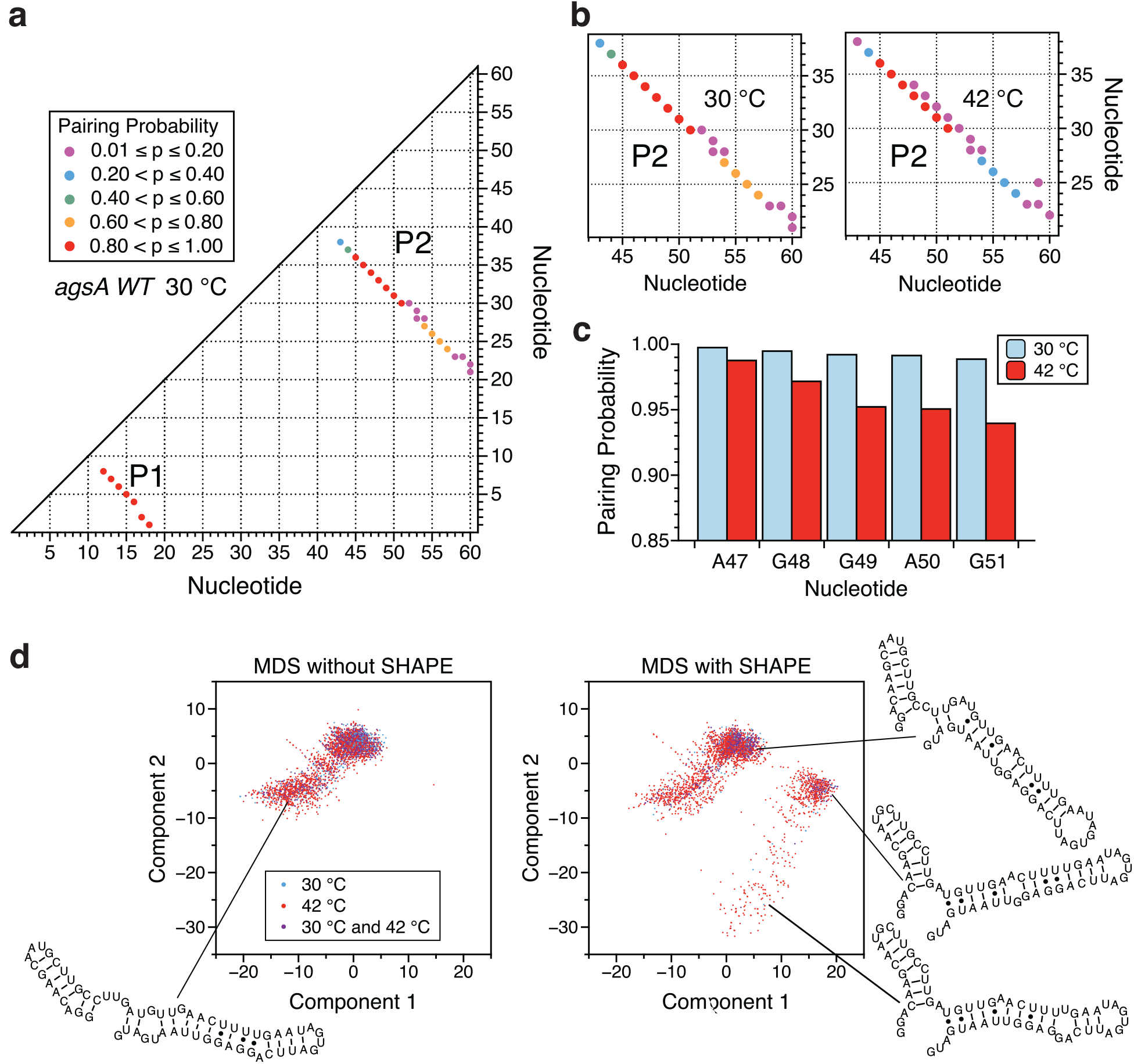
Ensemble visualization of the *agsA* WT thermometer modeled secondary structures at 30 °C and 42 °C. (**a**) Pairing-probability plot for the *agsA* WT thermometer at 30 °C. Dots correspond to base-pairs that occur in more than 1% of the predicted structural ensemble, as restrained by *in vivo* SHAPE-Seq data. Nucleotide numbers are plotted along the axes, with the x and y coordinates of a dot indicating the bases involved in the pair. Dot color corresponds to pairing probability, with red being the most likely and purple the least likely. (**b**) Close-up of in *vivo* SHAPE-Seq-restrained pairing-probability plot for the *agsA* WT thermometer hairpin 2 at 30 °C and 42 °C. Changes in pairing probability are indicative of the opening of the helix with increasing temperature. (**c**) The base-pairing probabilities for the first five nucleotides of the Shine-Dalgarno sequence as determined by the structural ensemble predictions restrained by *in vivo* SHAPE-Seq data, plotted at both 30 °C and 42 °C. (**d**) Multidimensional scaling (MDS) representations of the structural ensemble of the *agsA* WT thermometer at 30 °C and 42 °C, with and without *in vivo* SHAPE-Seq restraint of structural prediction. For each condition, 100,000 structures were sampled from the corresponding partition function and visualized in 2-dimensional space via MDS as described in the Experimental Procedures section. Representative structures shown.

## Results

### Characterization of in vivo structure/function of the agsA thermometer

As a first step, we characterized the function and structure of the *agsA* thermometer *in vivo*. To measure the thermometer’s temperature-sensitivity, we fused the *agsA* 5’ UTR directly to the super-folder green fluorescent protein (SFGFP) coding sequence and tested the construct’s heat-responsiveness *in vivo* in *Escherichia coli* (Figure 1). Alongside the wild-type (WT) *agsA* thermometer, we also tested a control 5’ UTR known to allow downstream translation at all temperatures and a pair of *agsA* mutants known to exhibit altered temperature responsiveness.^13^ Specifically, the G21C mutation is predicted to extend the *agsA* P2 helix to include the AUG start codon at its base, resulting in tighter repression of translation at 30 °C than the wild-type. The A29C mutation removes a predicted single base-pair internal hairpin loop, thus strengthening the P2 helix and preventing RBS access at both 30 °C and 42 °C (Figure 1A).

To test the responsiveness of the constructs to heat shock, we transformed them into *E. coli* and grew triplicate colonies overnight in liquid media. Duplicate subcultures of each colony were grown at 30 °C for 4 hours, and then one was subjected to a 30-minute heat shock at 42 °C while the other remained at 30 °C (Figure S2). From these cultures, we measured both fluorescent gene expression and structure of the *agsA* RNA with in-cell SHAPE-Seq.^21^

Gene expression was measured by determining fluorescence and optical density of the cultures using an optical plate-reader, which were used to calculate the normalized fluorescence units over optical density (FU/OD) values presented in Figure 1B. The control construct showed a slight increase in normalized fluorescence under heat shock, possibly corresponding to the faster production and maturation of SFGFP at the higher temperature. However compared to the control, the WT *agsA* variant showed clear activation in response to heat shock, with a 2.5-fold increase in normalized fluorescence over the 30 °C condition. In contrast, the G21C *agsA* variant gave lower overall levels of fluorescence but still showed increased gene expression in response to heat shock, while the A29C variant SFGFP expression levels were barely distinguishable above background at both 30 °C and 42 °C (Figure 1B). These results confirm the behavior of the *agsA* thermometer and its designed mutants in our expression configuration as originally reported in Walminghaus et al. using a different reporter system.^13^

Since mutations that stabilized the hairpin resulted in decreased gene expression, the results above support the model of thermometer regulation tied to hairpin melting at higher temperatures, though they do not detail the specific RNA structural changes that take place upon heat shock. To determine these structural changes *in vivo*, we probed the structure of the *agsA* 5’ UTR using in-cell SHAPE-Seq,^21,22^, in the same cell cultures used to make gene expression measurements. SHAPE-Seq is an RNA structure probing technique that uses a small chemical probe that covalently modifies RNA nucleotides in a structure-dependent fashion.^29^ The probe used here, 1M7 (1-methyl-7-nitroisatoic anhydride) also allows for in-cell probing of RNA structure.^21,22^ In general, the SHAPE family of probes preferentially react with unconstrained nucleotides like those in loops and single stranded regions, but do not frequently modify nucleotides constrained by base-pairing, stacking interactions, or ligand interactions.^31^ The locations of modifications are detected through reverse transcription and sequencing, allowing for the determination of the relative reactivities of the underlying RNA nucleotides to the SHAPE-Seq probe. High reactivities correspond to unpaired or unconstrained nucleotides, while low reactivities indicate base-pairing or constrained stacking interactions. These experimentally measured SHAPE-Seq reactivities can be incorporated into structural prediction programs as a pseudo-energy restraint that favors structures matching the observed reactivity patterns in order to arrive at highly accurate predictions of RNA secondary structure.^24–26^

We applied in-cell SHAPE-Seq to the *agsA* thermometer variants using reverse transcription primers that bound to the downstream coding sequence of SFGFP. Cultures were probed at 30 °C and 42 °C simultaneously with the bulk fluorescence assays presented in Figure 1. The resulting reactivity profiles (Figure 2a) were used to restrain RNA structural prediction (Figure 2b) using the RNAStructure software package^26^ to predict the minimum free energy (MFE) thermometer structures at each temperature. A comparison of the reactivity profiles and MFE folds between *agsA* variants and temperatures reveals a number of interesting differences.

For the wild-type *agsA* RNA, slight reactivity increases in the nucleotides comprising the base of the P2 helix under heat shock correspond to a predicted partial melting of the P2 hairpin at 42 °C. Though the reactivity changes are small, they correlate with the functional data showing increased gene expression at higher temperatures, potentially mediated by increased ribosome access to the Shine-Dalgarno region as the hairpin partially melts. While the G21C mutation restrained model does extend the P2 helix to contain the start codon at 30 °C, slight reactivity increases at 42 °C match a predicted expansion of the internal hairpin loop, correlating with the observed increase in fluorescence under heat shock conditions. Notably, the wild-type temperature transition was not correctly predicted by RNAStructure in the absence of SHAPE-Seq restraints (Figure S7); rather, the subtle reactivity changes measured by SHAPE-Seq influenced the folding algorithm enough to alter the predicted structures for the wild-type variant. In comparison, the A29C mutation that closes the inner hairpin loop creates a variant with strikingly low reactivities throughout the P2 helix and that displays no predicted structural change between 30 °C and 42 °C, retaining an extended hairpin form even under heat shock. This lack of hairpin opening corresponds to nearly undetectable levels of gene expression at both temperatures, bolstering the proposed mechanism of translational control through hairpin opening and ribosome access.

Despite the overall agreement with experimentally-informed predicted structural changes, it is worth noting that even at 42 °C, the MFE wild-type structure is still predicted to have near perfect base-pairing of the Shine-Dalgarno sequence due to the subtle nature of the observed reactivity changes. This suggests a more nuanced model of thermometer function than widespread structural changes from hairpin melting increasing ribosome access.^13^ Instead, the SHAPE-Seq data seem to indicate that only a small fraction of the RNA population is undergoing considerable structural changes. To examine this further, we used the RNAStructure `ProbabilityPlot` function to calculate the experimentally restrained ensemble base-pairing probabilities for the WT variant at 30 °C and 42 °C (Figure 3a and 3b). Points in these pairing-probability matrices reflect the probability that two bases will pair across the entire ensemble of structures, with helix elements showing up as diagonals in the matrix. Careful inspection of the probability matrices shows shifts toward lower probability base-pairing in the P2 stem at the higher temperature (Figure 3b), with Shine-Dalgarno base-pairing probabilities decreasing slightly (Figure 3c).

To obtain a more intuitive view of the population shift effect, we sampled structures from the experimentally restrained equilibrium partition function with RNAStructure’s `stochastic` method, and projected these structures onto a two-dimensional plane with multidimensional scaling (MDS) (Figure 3d). This analysis reveals that at 30 °C, the ensemble is dominated by a single cluster of similar closed structures, while at 42 °C a small fraction of the population shifts toward structures that have some bases unpaired, but only when experimental probing data are taken into account.

Consistent with these analyses, we observe only slight changes in the reactivity of the Shine-Dalgarno sequence (Figure 2a), with the experimentally restrained calculated ensemble-pairing probabilities shifting by just a few percentage points from approximately 99% pairing to 94-98% pairing at higher temperatures (Figure 3c). This subtle shift could correspond to a small population of transiently “breathing” hairpins. Intriguingly, this small ensemble shift in RBS accessibility is clearly enough to more than double the expression of the downstream reporter gene, demonstrating how subtle shifts in RNA structural ensembles can lead to large changes in functional output.

### Establishing the mechanism of the agsA thermometer’s functional response to temperature

Having observed *agsA* thermometer structure and function within the complex cellular environment where transcription and translation happen simultaneously, we next sought to use *in vitro* studies to decouple the two processes and better discern the underlying mechanism behind the *agsA* thermometer’s functional response to temperature. In particular, there is a possibility that temperature could influence the way the RNA molecule folds co-transcriptionally,^30^ which could then impact downstream translation of the mRNA. On the other hand, a simpler and purely post-transcriptional mechanism would rely on the ensemble of existing RNA folds adjusting to a temperature change, thus leading to changes in translation.

To distinguish between the two, we first tested the latter case by performing functional gene expression assays in the *in vitro* PURExpress system from purified *agsA* mRNAs. PURExpress is an *in vitro* protein synthesis system that can be used to express proteins from plasmid DNA or from purified mRNAs.^32^ We first established that the *agsA* thermometer variants gave the expected reporter output in PURExpress from plasmid input, allowing for both transcription and translation (Supplementary Figure S8). Next, we investigated whether we could observe the same functional response to temperature from purified *agsA* mRNAs. To do so, we tracked fluorescent reporter gene expression via plate reader for PURExpress reactions containing equal amounts of pre-transcribed, purified mRNAs for the control and *agsA* variants, thus eliminating any effect from co-transcriptional folding at a particular temperature. The resulting hour-long fluorescence traces (Figure 4a) recapitulated the function observed *in vivo*, demonstrating higher fluorescence relative to the control at 42 °C than 30 °C for the WT and G21C mutant and consistently low fluorescence for the A29C variant. The baseline increase in gene expression of the control from 30 °C to 42 °C is higher than seen *in vivo*, which could be due partly to the longer duration of the heat shock and partly to the magnifying effect of the high-yield PURExpress system. Figure 4b presents the measured rates of fluorescence increase during linear growth, reflecting the same trends observed in overall fluorescence. While these experiments cannot rule out cotranscriptional effects completely, the significant temperature responsiveness of the pre-transcribed *agsA* construct mRNAs demonstrates that this RNA thermometer can function through thermodynamic shifts in the RNA fold ensemble as was previously suggested by Waldminghaus et al.^13^

**Figure 4.**
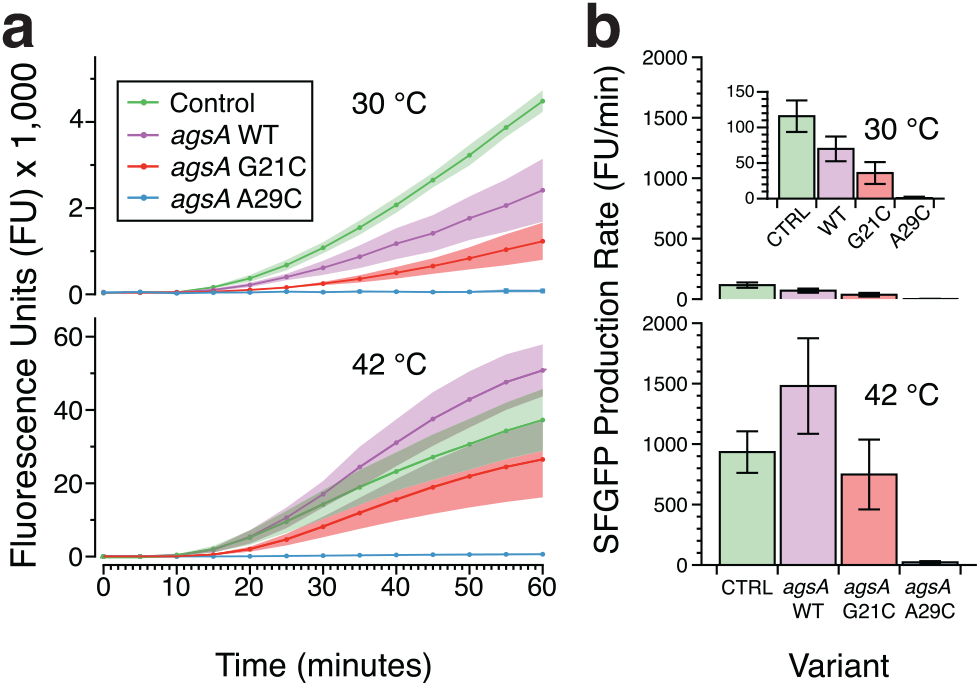
Testing *agsA* thermometer function in a cell-free protein synthesis system using purified pre-transcribed mRNA. (**a**) Fluorescence trajectories over time for translation of SFGFP from *agsA* constructs and control mRNA in the PURExpress protein synthesis system at 30 °C and 42 °C. Shading represents standard deviation over three replicates. (**b**) SFGFP production rates, calculated from the trajectories in (a), during the linear synthesis regime for *agsA* constructs and control at 30 °C (45-50 minutes) and 42 °C (30-35 minutes), with error bars representing standard deviation.

### Investigating agsA thermometer folding in vitro using SHAPE-Seq

Having confirmed the basic mechanism of *agsA* thermometer function, we set about investigating *agsA* thermometer folding *in vitro* to explore whether we could detect a stronger signal for structural switching in the absence of other cellular factors. Performing SHAPE-Seq *in vitro* in the PURExpress cell-free system without ribosomes on *agsA* WT mRNA at 30 °C and 42 °C produced the reactivity plots presented in Figure 5a. Quite notably, the reactivity profiles at both temperatures are markedly similar to those measured *in vivo*, with only a handful of nucleotides showing statistically significant differences in reactivity. When the *in vitro* reactivities are used to restrain the RNA structural prediction, the resulting structural predictions (Figure 5b) parallel those seen in under *in vivo* conditions, but with two key differences: at 30 °C the internal loop in the P2 helix is predicted to be one base-pair smaller *in vitro*, and at 42 °C the P2 stem loop is predicted to be two base-pairs larger *in vitro*. Similar patterns in reactivity and structure were seen for *in vitro* experiments conducted in buffer as opposed to PURExpress, demonstrating a stable pattern of helix formation with slight variations in loop size. (See Figures S9 and S10.) Overall, the close agreement between *in vivo* reactivities and *in vitro* reactivities in the absence of active translation suggests that the cellular translation machinery is exploiting subtle changes in RNA helix opening to enact large changes in gene expression.

**Figure 5.**
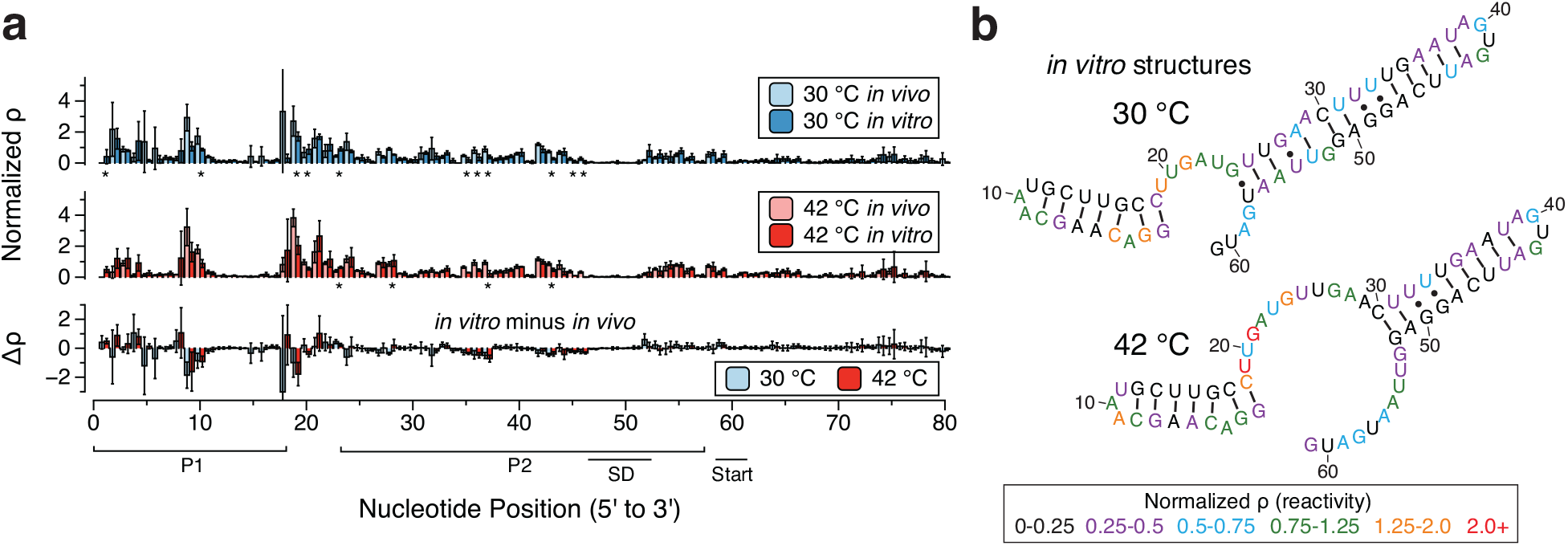
SHAPE-Seq characterization of the *agsA* thermometer *in vitro* in PURExpress. (**a**) SHAPE-Seq normalized reactivity plots for *agsA* WT mRNA from triplicate experiments *in vitro* performed on mRNAs at 30 °C and 42 °C, graphed alongside *in vivo* reactivities from Figure 2a for comparison. Error bars represent standard deviations over replicates. Stars denote statistically significant differences between *in vivo* and *in vitro* as determined by heteroscedastic Welch’s T-tests, while the difference plot shows the magnitude of reactivity change *(in vitro* reactitivies minus *in vivo* reactivities). (**b**) Experimentally-informed minimum free energy (MFE) RNA structures as predicted using the RNAStructure `Fold` program with the *in vitro* reactivity values from (a) taken as pseudoenergy folding restraints. Nucleotides are color-coded according to normalized SHAPE reactivity (ϱ), with red indicating high reactivity and black low reactivity.

### Determining the effect of translation and ribosome binding on agsA RNA structure

We next sought to uncover the influence of ribosome binding on *agsA* thermometer structure. Specifically, we performed additional SHAPE-Seq experiments *in vitro* in PURExpress involving both active translation and stalled translation initiation complexes (TIC) to determine the reactivity changes caused by ribosome binding and active translation in the clearly defined *in vitro* system (Figure 6). For the active translation experiments, we probed PURExpress reactions in the same conditions used for functional studies. The reactivities in the *agsA* 5’UTR are relatively similar in the no ribosomes and active ribosome conditions (Figure 6b), with slight reactivity increases in the P2 helix. Interestingly, we also observe reactivity differences within the coding sequence, where reactivities are lower with active ribosomes, most likely due to actively translating ribosomes providing protection from SHAPE modification.

**Figure 6.**
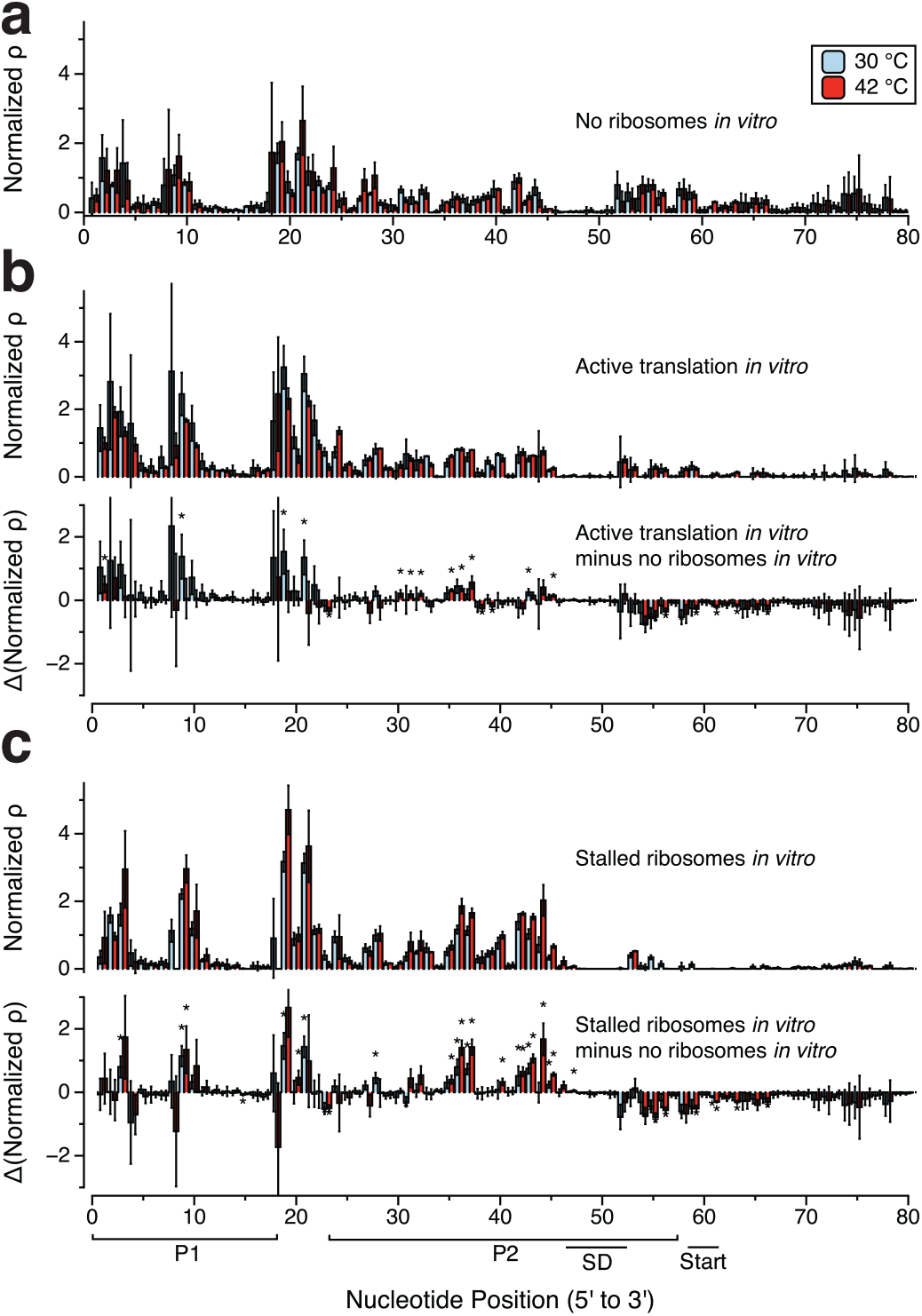
Investigating the effect of translation and ribosome occupancy on *agsA* WT structure and reactivity. (**a**) *agsA* WT SHAPE-Seq reactivity plot for triplicate experiments carried out *in vitro* in PURExpress without ribosomes. (**b**) *agsA* WT SHAPE-Seq reactivity plot in PURExpress with active translation and difference plot of reactivities showing the change from the no ribosome case in part (a). (**c**) *agsA* WT SHAPE-Seq reactivity plot in PURExpress with stalled translation initiation complexes generated by supplying only methionine and difference plot of reactivities showing the change from the no ribosome case in part (a). Error bars represent standard deviations over replicates. Stars represent statistical significance as in Figure 5.

To probe stalled TIC complexes, we probed PURExpress reactions assembled lacking all amino acids except for methionine, thus causing the translation initiation complex to stall after the addition of the first amino acid. As in the active ribosome experiment, there is a stark reactivity drop directly downstream of the ribosome binding site, likely corresponding to the ribosome footprint blocking SHAPE modification (Figure 6c). The large size of the region of reactivity drop extending ~20 nt beyond the start site is consistent with previous observations of stalled ribosomes in the presence of tRNA(fMet).^33^ Even more interestingly, many of the nucleotides that formed the P2 helix, on the 5’ side and the upper portion of the 3’ side, undergo large increases in reactivity in the stalled TIC complex condition. This suggests increased nucleotide flexibility brought about by the dissolution of the P2 helix in concert with ribosome binding. Larger changes in reactivity at 42 °C than 30 °C suggest greater ribosome access at the higher temperature, matching the functional data seen *in vivo* and *in vitro*. However, given that the ribosomes were supplied in excess to the mRNA (8:1 ratio) and allowed 20 minutes to reach binding equilibrium, it is unsurprising that the reactivities indicate ribosome binding to a large portion of the mRNA ensemble under both temperature conditions. The presence of background gene expression at 30 °C in functional testing indicates some level of ribosome access even at the lower temperature, most likely linked to a small fraction of the ensemble undergoing transient hairpin breathing that allows for ribosome capture. Overall this data suggests an active role for ribosome binding in mediating additional RNA structural changes that facilitate *agsA* thermometer function.

## Discussion

In this work, we have combined *in vivo* and *in vitro* structure-function probing experiments to uncover detailed mechanistic aspects of how the *agsA* RNA thermometer activates gene expression in response to heat shock. Specifically, these experiments represent the first *in vivo* RNA structure probing of an RNA thermometer and nicely complement previous *in vitro* structural studies of fourU thermometers.^13,19,20^ Overall, our *in vivo* SHAPE-Seq data revealed only small structural changes in the *agsA* thermometer with an increase in temperature, indicating subtle shifts in the underlying RNA structural ensemble upon shifting to higher temperatures. This is consistent with previous *in vitro* NMR structural studies that showed gradual base-pair dissociation at higher temperatures.^19^ Thus it appears that temperature shifts do not cause large-scale structural rearrangement in this simple hairpin. However, our subsequent *in vitro* functional experiments showed that these subtle equilibrium-based post-transcriptional changes in RNA structure do lead to temperature-dependent control of gene expression. This is despite the lack of major temperature-dependent structural differences in the Shine-Dalgarno region, suggested as the mechanism behind *agsA* thermometer function.^13^

Further exploration of *agsA* thermometer structure with active or stalled ribosomes did reveal an important mechanistic insight: ribosomes can exploit small changes in the structural ensemble to increase translation of mRNAs at elevated temperatures. Specifically, we observed an increase in reactivities in the region paired to the Shine-Dalgarno site when stalled ribosomes were present, suggesting that ribosome binding can unwind the *agsA* thermometer’s P2 helix to facilitate translation initiation. This happens during heat shock conditions and to a lesser degree at lower temperatures, as evidenced by the changes in reactivities surrounding the ribosome binding site at 30 °C in Figure 6c, as well as the background level of gene expression detectable at 30 °C. These observations reinforce the idea that the ribosome is able to take advantage of very subtle shifts in RNA structure, even when the majority of the population of RNA thermometer hairpins is well-folded.

In conclusion, we have demonstrated strong evidence that subtle differences in RNA structures mediated by temperature changes can be exploited by ribosomes to give rise to large changes in gene expression. It is interesting to speculate whether this mechanism could be important for the general control of gene expression in a temperature-dependent manner. Not only does the elegantly simple hairpin fluctuation coupled to ribosome-mediated melting mechanism behind *agsA* thermo-sensing offer a potential template for the future design of synthetic RNA thermometers,^34–36^ but the combination of SHAPE-Seq with *in vitro* mechanistic studies shows how *in vivo* probing can be used to validate the structural mechanism behind regulatory RNAs.

## Acknowledgements

We acknowledge Matt Verosloff for conducting in-house replication experiments, James Chappell and Kyle Watters for fruitful discussions about experimental design, Angela Yu for assistance with computational analysis, and Paul Carlson for providing the control construct. We also thank Franz Narberhaus of Ruhr-Universität Bochum for providing *agsA* plasmids. This work was supported by the National Science Foundation Graduate Research Fellowship Program [grant number DGE-1144153 to SM] and a New Innovator Award through the National Institute of General Medical Sciences of the National Institutes of Health [grant number 1DP2GM110838 to JBL]. The content is solely the responsibility of the authors and does not necessarily represent the official views of the National Institutes of Health.

Supporting Information Available: Supplementary Figures S1-S10 and Supplementary Table S1.

The authors declare no competing financial interests.

## References

(1) Kortmann, J., and Narberhaus, F. (2012) Bacterial RNA thermometers: molecular zippers and switches. Nat. Rev. Microbiol. 10, 255–265.

(2) Konkel, M. E., and Tilly, K. (2000) Temperature-regulated expression of bacterial virulence genes. Microbes Infect. 2, 157–166.

(3) Shapiro, R. S., and Cowen, L. E. (2012) Thermal control of microbial development and virulence: Molecular mechanisms of microbial temperature sensing. MBio 3, 1–6.

(4) Narberhaus, F. (2010) Translational control of bacterial heat shock and virulence genes by temperature-sensing mRNAs. RNA Biol. 7, 84–89.

(5) de Smit, M. H., and van Duin, J. (1990) Secondary structure of the ribosome binding site determines translational efficiency: a quantitative analysis. Proc. Natl. Acad. Sci. 87, 7668–72.

(6) Waldminghaus, T., Fippinger, A., Alfsmann, J., and Narberhaus, F. (2005) RNA thermometers are common in alpha- and gamma-proteobacteria. Biol. Chem. 386, 1279–86.

(7) Waldminghaus, T., Gaubig, L. C., Klinkert, B., and Narberhaus, F. (2009) The *Escherichia coli ibpA* thermometer is comprised of stable and unstable structural elements. RNA Biol. 6, 455–463.

(8) Chowdhury, S., Maris, C., Allain, F. H.-T., and Narberhaus, F. (2006) Molecular basis for temperature sensing by an RNA thermometer. EMBO J. 25, 2487–97.

(9) Morita, M., Kanemori, M., Yanagi, H., and Yura, T. (1999) Heat-induced synthesis of σ32 in Escherichia coli: structural and functional dissection of rpoH mRNA secondary structure. J. Bacteriol. 181, 401–10.

(10) Morita, M. T., Tanaka, Y., Kodama, T. S., Kyogoku, Y., Yanagi, H., and Yura, T. (1999) Translational induction of heat shock transcription factor σ32: evidence for a built-in RNA thermosensor. Genes Dev. 13, 655–65.

(11) Wagner, D., Rinnenthal, J., Narberhaus, F., and Schwalbe, H. (2015) Mechanistic insights into temperature-dependent regulation of the simple cyanobacterial hsp17 RNA thermometer at base-pair resolution. Nucleic Acids Res. 43, 5572–5585.

(12) Kortmann, J., Sczodrok, S., Rinnenthal, J., Schwalbe, H., and Narberhaus, F. (2011) Translation on demand by a simple RNA-based thermosensor. Nucleic Acids Res. 39, 2855–68.

(13) Waldminghaus, T., Heidrich, N., Brantl, S., and Narberhaus, F. (2007) FourU: a novel type of RNA thermometer in Salmonella. Mol. Microbiol. 65, 413–24.

(14) Tomoyasu, T., Takaya, A., Sasaki, T., Nagase, T., Kikuno, R., Morioka, M., and Yamamoto, T. (2003) A New Heat Shock Gene, agsA, Which Encodes a Small Chaperone Involved in Suppressing Protein Aggregation in Salmonella enterica Serovar Typhimurium. J. Bacteriol. 185, 6331–6339.

(15) Hoe, N. P., and Goguen, J. D. (1993) Temperature sensing in *Yersinia pestis*: translation of the LcrF activator protein is thermally regulated. J. Bacteriol. 175, 7901–7909.

(16) Weber, G. G., Kortmann, J., Narberhaus, F., and Klose, K. E. (2014) RNA thermometer controls temperature-dependent virulence factor expression in Vibrio cholerae. Proc. Natl. Acad. Sci. 111, 14241–14246.

(17) Johansson, J., Mandin, P., Renzoni, A., Chiaruttini, C., Springer, M., and Cossart, P. (2002) An RNA thermosensor controls expression of virulence genes in Listeria monocytogenes. Cell 110, 551–561.

(18) Loh, E., Dussurget, O., Gripenland, J., Vaitkevicius, K., Tiensuu, T., Mandin, P., Repoila, F., Buchrieser, C., Cossart, P., and Johansson, J. (2009) A trans-Acting Riboswitch Controls Expression of the Virulence Regulator PrfA in Listeria monocytogenes. Cell 139, 770–779.

(19) Rinnenthal, J., Klinkert, B., Narberhaus, F., and Schwalbe, H. (2010) Direct observation of the temperature-induced melting process of the Salmonella fourU RNA thermometer at base-pair resolution. Nucleic Acids Res. 38, 3834–47.

(20) Rinnenthal, J., Klinkert, B., Narberhaus, F., and Schwalbe, H. (2011) Modulation of the stability of the Salmonella fourU-type RNA thermometer. Nucleic Acids Res. 39, 8258–8270.

(21) Watters, K. E., Abbott, T. R., and Lucks, J. B. (2015) Simultaneous characterization of cellular RNA structure and function with in-cell SHAPE-Seq. Nucleic Acids Res. 44.

(22) Watters, K. E., Yu, A. M., Strobel, E. J., Settle, A. H., and Lucks, J. B. (2016) Characterizing RNA structures in vitro and in vivo with selective 2’-hydroxyl acylation analyzed by primer extension sequencing (SHAPE-Seq). Methods 103, 34–48.

(23) Aviran, S., Trapnell, C., Lucks, J. B., Mortimer, S. A., Luo, S., Schroth, G. P., Doudna, J. A., Arkin, A. P., and Pachter, L. (2011) Modeling and automation of sequencing-based characterization of RNA structure. Proc. Natl. Acad. Sci. 108, 11069–74.

(24) Loughrey, D., Watters, K. E., Settle, A. H., and Lucks, J. B. (2014) SHAPE-Seq 2.0: systematic optimization and extension of high-throughput chemical probing of RNA secondary structure with next generation sequencing. Nucleic Acids Res. 42, 1–10.

(25) Low, J. T., and Weeks, K. M. (2010) SHAPE-directed RNA secondary structure prediction. Methods 52, 150–8.

(26) Reuter, J. S., and Mathews, D. H. (2010) RNAstructure: software for RNA secondary structure prediction and analysis. BMC Bioinformatics 11, 129.

(27) Hofacker, I. L. (2003) Vienna RNA secondary structure server. Nucleic Acids Res. 31, 3429–3431.

(28) Kutchko, K. M., Sanders, W., Ziehr, B., Phillips, G., Solem, A., Halvorsen, M., Weeks, K. M., Moorman, N., and Laederach, A. (2015) Multiple conformations are a conserved and regulatory feature of the RB1 5’ UTR. RNA 21, 1274–85.

(29) Lucks, J. B., Mortimer, S. A., Trapnell, C., Luo, S., Aviran, S., Schroth, G. P., Pachter, L., Doudna, J. A., and Arkin, A. P. (2011) Multiplexed RNA structure characterization with selective 2’-hydroxyl acylation analyzed by primer extension sequencing (SHAPE-Seq). Proc. Natl. Acad. Sci. 108, 11063–8.

(30) Watters, K. E., Strobel, E. J., Yu, A. M., Lis, J. T., and Lucks, J. B. (2016) Cotranscriptional folding of a riboswitch at nucleotide resolution. Nat. Struct. Mol. Biol. 23, 1124–1131.

(31) Bindewald, E., Wendeler, M., Legiewicz, M., Bona, M. K., Wang, Y., Pritt, M. J., Le Grice, S. F. J., and Shapiro, B. a. (2011) Correlating SHAPE signatures with three-dimensional RNA structures. RNA 17, 1688–1696.

(32) Tuckey, C., Asahara, H., Zhou, Y., and Chong, S. (2014) Protein synthesis using a reconstituted cell-free system. Curr. Protoc. Mol. Biol. 2014, 16.31.1–16.31.22.

(33) Hüttenhofer, A., and Noller, H. F. (1994) Footprinting mRNA-ribosome complexes with chemical probes. EMBO J. 13, 3892–901.

(34) Hoynes-O’Connor, A., Hinman, K., Kirchner, L., and Moon, T. S. (2015) De novo design of heat-repressible RNA thermosensors in E. coli. Nucleic Acids Res. 43, 6166–6179.

(35) Neupert, J., and Bock, R. (2009) Designing and using synthetic RNA thermometers for temperature-controlled gene expression in bacteria. Nat. Protoc. 4, 1262–1273.

(36) Waldminghaus, T., Kortmann, J., Gesing, S., and Narberhaus, F. (2008) Generation of synthetic RNA-based thermosensors. Biol. Chem. 389, 1319–26.

